# Refining bias correction in genome-wide association analyses of case-control studies

**DOI:** 10.1101/2025.09.01.673522

**Authors:** Behrooz Darbani, Ole Birger Vesterager Pedersen, Sisse Rye Ostrowski, Qihua Tan, Vibeke Andersen

## Abstract

Genome-wide association studies are vulnerable to confounding factors. This study provides evidence-based guidance for minimizing bias associated with genetic relatedness, SNP-specific non-additive allelic interactions, predisposed genotypes among controls, and multi-allelic polymorphism in case-control studies. The analyses demonstrated that genetic similarity within case or control groups introduces experimental bias, whereas genetic relatedness across case-control samples reduces this bias. These findings establish a general framework that can filter genetically related sub-communities or paired samples, whilst preserving maximal statistical power with minimal false-positive rates. Moreover, the skewed odds ratios resulting from predisposed genotypes among controls underscored the importance of age-related filtering to minimize this confounding effect. To ensure accurate genetic estimates, such as polygenic risk scores, the identification of SNP-specific allelic interaction models was also emphasized in case-control studies, contingent on normalization for within-population differences in genotype frequencies. Here, we introduce the Allelic-effect aware Case-Control GWAS (AlleliC-GWAS) tool for identifying SNP-specific allelic effects on binary traits. Finally, we recommend a strategy to accurately capture genetic effects at multi-allelic genomic positions.

## Background

Genome-wide association studies are designed to identify genetic variants that contribute to trait variation. The advent of advanced sequencing technologies marked a paradigm shift from traditional molecular markers, such as Restriction Fragment Length Polymorphism (RFLP), to whole genome sequences [1–4]. This transition facilitated the identification of millions of polymorphic genomic loci, driving a surge in genome-wide association studies. Despite recent advancements in genomic and genotypic coverage, i.e., analyzing large cohorts using millions of single-nucleotide polymorphisms (SNPs) [5–11], methodological considerations in genetic data analysis are indispensable to reduce bias and thereby, the number of false-positive and negative SNP hits. This study outlines strategies to maximize statistical power for the discovery of causal SNPs while minimizing spurious associations and improving the precision of genetic estimates.

Like population-level genetic differences [12], individual-level genetic similarities can bias genetic assessments. Genetic relatednesses, such as pedigree and cryptic similarities, have been identified as major confounders in genome-wide association studies, leading to inflated type I errors [13–20]. Although mixed models [18,20–27] have been introduced to control for sample relatedness, sample filtering by retaining one individual per related group of individuals has been the most common approach [28–35]. In contrast, family-based association studies can offer greater statistical power than case-control association studies of genetically unrelated individuals [36–41]. However, the advantage of family-based association studies can be easily undermined by the typically small sample sizes inherent to such designs. In this study, we explored optimal strategies for filtering genetically related samples to control the incidence of false SNP hits effectively.

The manifestation of specific traits, such as the onset of some diseases, occurs at different ages throughout the human lifespan. This can introduce bias in case-control genome-wide studies through recruiting individuals mistakenly classified as controls, e.g., healthy individuals who do not have a given chronic disease but carry causal genetic elements that will express phenotypically at higher ages. This impurity among controls, i.e., the presence of predisposed genotypes, can lead to inaccurate genetic measures such as underestimated odds ratios. The presence of predisposed genotypes may arise irrespective of recruitment scheme, i.e., under both active participation (non-random) and passive participation strategies. Accordingly, methods that address active participation-mediated volunteer bias [34], also known as participation bias [42], do not interfere with bias mediated by predisposed genotypes. Furthermore, the GWAS-by-proxy (GWAX) methodology [43], which uses parental diagnostic history, can potentially introduce biases due to uncertainty in phenotypic concordance between parents and offspring. Accordingly, the application of GWAX has been found to generate misleading outcomes [44]. Here, we propose a filtering strategy to reduce impurity among control samples within cohorts.

Another important aspect that can easily influence the accuracy of genetic measures is the type of interaction between the reference and alternative alleles at a given polymorphic genomic position. These interactions should be distinguished from epistatic interactions among SNPs [45,46]. Recent studies have reported dominance effects in mammals [47] and humans [48]. In contrast to pervasive non-linear interactions among genes or SNPs [46,49], there has been a widespread assumption that SNP-specific non-additive allelic effects are negligible [50]. Nevertheless, this assumption does not rest on any inherent genetic or biological principles. Sample-size enlargement is required for capturing dominance effects efficiently at the levels observed for additive effects [51]. As illustrated, compared with the observed allelic effects, the estimated small non-additive allelic effects may be attributable to methodological limitations and artefacts [52]. Here, we highlight and adjust the application of recently introduced observed-level allelic effects [52] to efficiently capture additive and non-additive allelic effects, including dominance, overdominance, and heterosis-like effects, in case-control studies. We also shed light on the importance of normalizing differences in genotype frequencies to accurately capture additive and non-additive effects in case-control studies, such as those with binary phenotypes. We finally provide a brief methodological discussion on extracting the effect of each alternative allele at multi-allelic genomic positions. Each allele at a multi-allelic site is often encoded relative to all other alleles at that specific genomic position, which can hinder the accurate estimation of allele-specific genetic effects.

## Results

### Genetic relatedness and sample filtering

Within-cohort genetic relatedness is recognized as a major confounding factor, leading to inflated type I error rates in genome-wide association studies [13–20]. It has been a common strategy to avoid erroneous association outcomes by genetic relationship inference involving control and case samples, i.e., filtering out individuals with second-degree or higher levels of genetic relatedness and retaining only one individual per group of related individuals, with priority given to the case samples over controls [28–35]. As our results also prove (Figure 1-3), we must note that the confounding effects of relatedness are relevant only for the case-samples or control-samples specific relatedness, and not for relatedness across case and control groups. Sample-relatedness confined to either the case or control group can artificially inflate the frequency of unassociated alleles within the respective group, thereby generating spurious association signals. In contrast, the relatedness across case-control samples provides an even distribution of unassociated alleles between case and control groups; statistically significant association signals from unassociated SNPs are a common problem arising from low genotypic coverage. Therefore, genetic relatedness across case-control samples not only does not lead to false positives but also reduces them, mimicking the effect of family-based genetic studies. This mimicked effect resembles the superiority, i.e., higher statistical power and accuracy coupled with minimaized false-positive rates, reported for family-based genetic studies [36–41,53], assuming equivalent sample sizes.

**Figure 1.**
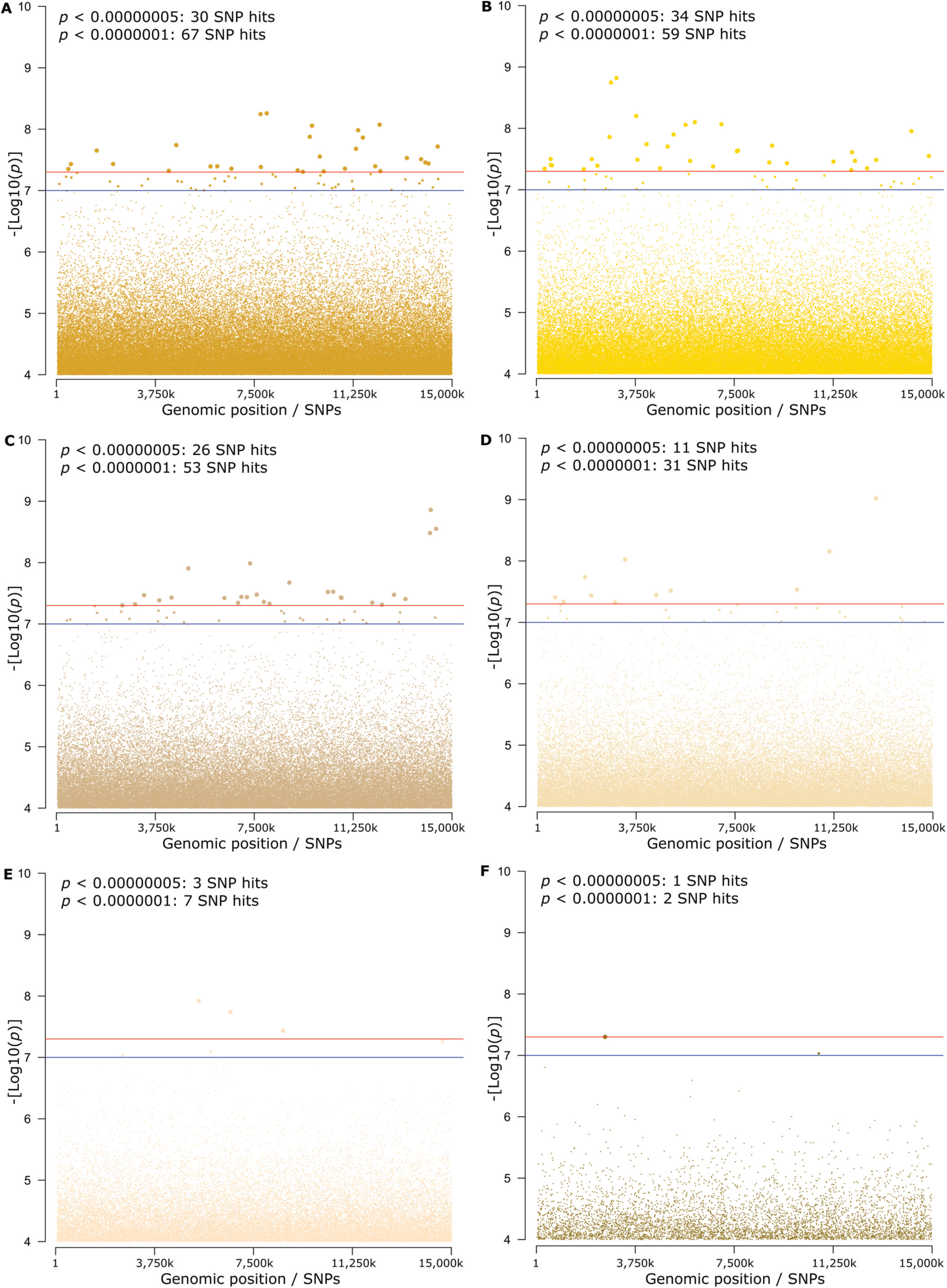
Genetic relatedness across case-control samples reduces random associations, i.e., false positive rates. Each Manhattan plot is illustrating the outputs from 60 independent replicates of case-control association analyses. The analyses were based on simulated data including 15 million SNPs (MAF ≥ 0.01) for 10,000 genetically unrelated case samples (genetic similarity ≤ 0.3%) and 10,000 genetically unrelated controls samples (genetic similarity ≤ 0.3%). The decreased number of random association signals in these datasets reflect reduced false positive rates resulting from random allocation of case and control samples. A) Genetic similarities between case and control samples were ≤ 0.3%, except for 100 control samples each sharing between 24.8%–25.1% genetic similarity with an individual case samples. B) Genetic similarities between case and control samples were ≤ 0.3%, except for 1,000 control samples each sharing between 24.8%–25.1% genetic similarity with an individual case samples. C) Genetic similarities between case and control samples were ≤ 0.3%, except for 2,000 control samples each sharing between 24.8%–25.1% genetic similarity with an individual case samples. D) Genetic similarities between case and control samples were ≤ 0.3%, except for 2,500 control samples each sharing between 24.8%–25.1% genetic similarity with an individual case samples. E) Genetic similarities between case and control samples were ≤ 0.3%, except for 5,000 control samples each sharing 24.8%–25.1% genetic similarity with an individual case samples. F) Genetic similarities between case and control samples were 24.8%–25.1%. MAF: Minor allele frequency.

**Figure 2.**
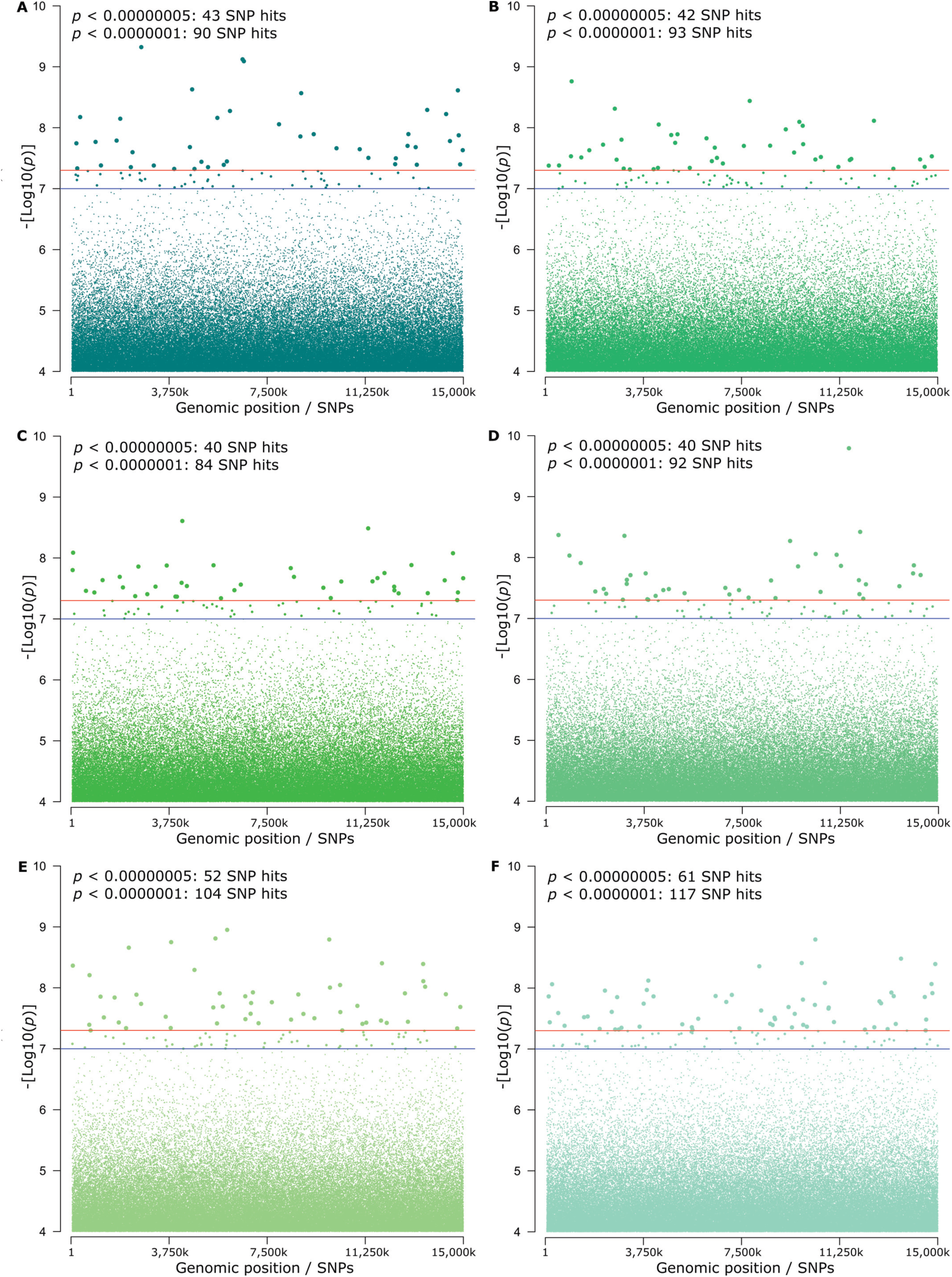
Random associations across varying numbers of genetically related, case- or control-specific paired samples. Each Manhattan plot is illustrating the outputs from 60 independent replicates of case-control association analyses. The analyses were based on simulated data including 15 million SNPs (MAF ≥ 0.01) and 20,000 samples; 10,000 case samples which were genetically unrelated, i.e., genetic similarity ≤ 0.3%, to 10,000 controls samples. The elevated numbers of random association signals in these datasets reflect increased false positive rates resulting from random allocation of case and control samples. A) Analyzing genetically unrelated samples. Genetic similarities among cases, controls, and case-control samples were ≤ 0.3%. B) Analyzing genetically unrelated samples. Genetic similarities among cases, controls, and case-control samples were ≤ 0.3%, except 50 paired samples from the control group exhibiting 24.8%–25.1% genetic similarity between paired samples. C) Analyzing genetically unrelated samples. Genetic similarities among cases, controls, and case-control samples were ≤ 0.3%, except 100 paired samples from the control group exhibiting 24.8%–25.1% genetic similarity between paired samples. D) Analyzing genetically unrelated samples. Genetic similarities among cases, controls, and case-control samples were ≤ 0.3%, except 250 paired samples from the control group exhibiting 24.8%–25.1% genetic similarity between paired samples. E) Analyzing genetically unrelated samples. Genetic similarities among cases, controls, and case-control samples were ≤ 0.3%, except 500 paired samples from the control group exhibiting 24.8%–25.1% genetic similarity between paired samples. F) Analyzing genetically unrelated samples. Genetic similarities among cases, controls, and case-control samples were ≤ 0.3%, except 1,000 paired samples from the control group exhibiting 24.8%–25.1% genetic similarity between paired samples. MAF: Minor allele frequency.

**Figure 3.**
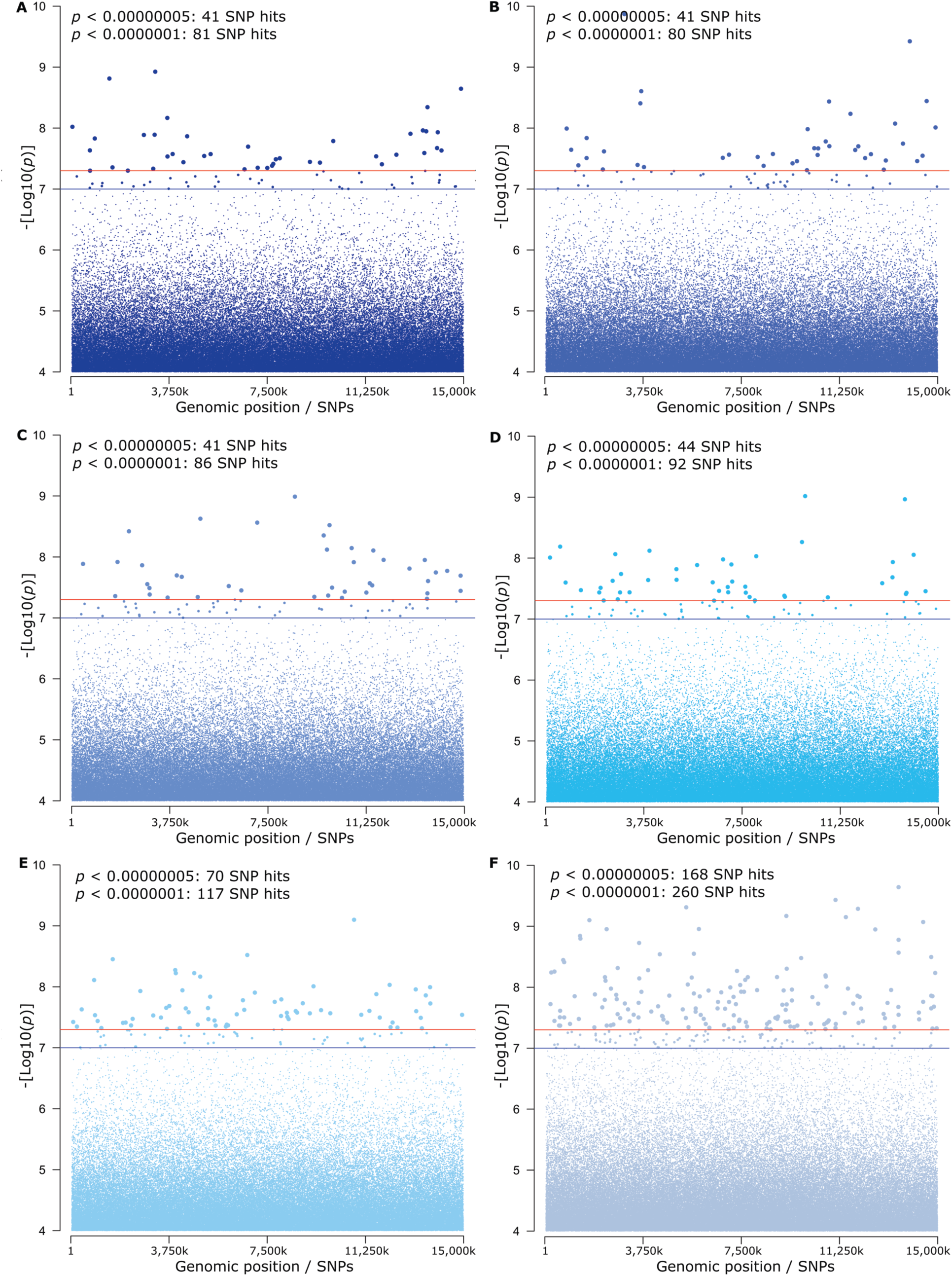
Random associations across varying sizes of genetically related case- or control-specific sub-communities. Each Manhattan plot is illustrating the outputs from 60 independent replicates of case-control association analyses. The analyses were based on simulated data including 15 million SNPs (MAF ≥ 0.01) and 20,000 samples; 10,000 case samples which were genetically unrelated, i.e., genetic similarity ≤ 0.3%, to 10,000 controls samples. The elevated numbers of random association signals in these datasets, relative to the dataset without relatedness (Figure 2A), reflect increased false positive rates resulting from random allocation of case and control samples. A) Analyzing genetically unrelated samples. Genetic similarities among cases, controls, and case-control samples were ≤ 0.3%, except for a sub-community of 10 control samples, each sharing 24.8% to 25.1% genetic similarity with all other samples within the sub-community. B) Analyzing genetically unrelated samples. Genetic similarities among cases, controls, and case-control samples were ≤ 0.3%, except for a sub-community of 15 control samples, each sharing 24.8% to 25.1% genetic similarity with all other samples within the sub-community. C) Analyzing genetically unrelated samples. Genetic similarities among cases, controls, and case-control samples were ≤ 0.3%, except for a sub-community of 20 control samples, each sharing 24.8% to 25.1% genetic similarity with all other samples within the sub-community. D) Analyzing genetically unrelated samples. Genetic similarities among cases, controls, and case-control samples were ≤ 0.3%, except for a sub-community of 25 control samples, each sharing 24.8% to 25.1% genetic similarity with all other samples within the sub-community. E) Analyzing genetically unrelated samples. Genetic similarities among cases, controls, and case-control samples were ≤ 0.3%, except for a sub-community of 35 control samples, each sharing 24.8% to 25.1% genetic similarity with all other samples within the sub-community. F) Analyzing genetically unrelated samples. Genetic similarities among cases, controls, and case-control samples were ≤ 0.3%, except for a sub-community of 50 control samples, each sharing 24.8% to 25.1% genetic similarity with all other samples within the sub-community. MAF: Minor allele frequency.

Figures 1-3 illustrate the advantage of genetic relatedness across case-control samples, in contrast to the disadvantages of case-samples specific or control-samples specific genetic relatedness in genome-wide association studies. Our results demonstrate the usefulness of relatedness between case and control samples for reducing false positive rates (Figure 1). Interestingly, a subset of 100 genetically related samples across case and control groups, accounting for just 1% of the cohort, exhibited a lower false-positive rate (Figure 1A, 2A). A sharp reduction in false positive rate was also observed in the presence of genetic relatedness across case and control groups, including all samples (Figure 1F, 2A). On the other hand, including ≥ 500 case or control group-specific genetically related paired samples, i.e., comprising ≥ 10% of that group, did raise the false-positive rate (Figure 2). Small genetically related sub-communities containing up to 25 related samples also did not elevate the false-positive rate (Figure 3). Taken together, these results provide a general guidance to handle the sample relatedness within cohorts via sample filtering: i) sample relatedness across case-control groups reduces false positive rates and thus obviates the need for filtering; ii) genetically related small sub-communities, e.g., sub-communities constituting 0.25% of controls or cases, do not lead to false positive SNP hits; and, iii) genetically related sample pairs confined to either the case or control group can lead to false-positive associations and warrant filtering when they constitute ≥ 10% of their respective group, i.e., case or control. For these simulations, we used a relatively high level of relatedness, i.e., 25% genetic similarity, among all the related samples, and thereby, the mentioned thresholds of 0.25% and 10% are quite stringent; higher threshold levels can be considered acceptable in many studies having lower levels of genetic relatedness among individuals.

Sample filtering has been the most common approach for controlling relatedness [28–35], and one of the primary goals of this study was to establish an efficient sample filtering strategy. For the purpose of comparison, we also performed analyses using the mixed-model fastGWA-GLMM [23] to control for relatedness without the need for sample filtering. As expected, the mixed-model results were comparable to those from logistic regression when analyzing the dataset with no relatedness (Figure 2A, S1A) or with relatedness across case-control groups (Figure 1F, S1B). The analyses by fastGWA-GLMM also demonstrated effective control of relatedness without sample filtering when we examined the previously described cohort, which contained a genetically related sub-community of 50 related samples within either the control or case group (Figure S1A,C). It is important to note that when mixed models are used to account for sample-relatedness, scenarios with case-specific or control-specific relatedness are also expected to yield similar numbers of random association hits as the corresponding scenarios lacking relatedness. We nevertheless observed an elevated false-negative risk when applying the mixed model to examine the previously described cohort with 1,000 genetically related paired samples within either the control or case group (Figure S1D). Specifically, the number of random associations with *p*-values less than 10^-8^ decreased from 90 in the scenario without related samples (Figure 2A and Figure S1A) to 58 in the scenario with 1,000 genetically related paired samples (Figure S1D), despite no underlying reason, i.e., without any related samples being shared between the case and control groups. We additionally detected a markedly increased number of random associations upon applying the mixed model following sample filtering (Figure S2). These analyses indicate that there is no single prescribed approach for choosing between sample filtering and mixed models for controlling relatedness. When applying the commonly used approach of sample filtering [28–35], our study recommends avoiding the removal of related samples across case and control groups.

### The effect of predisposed genotypes, i.e., individuals among controls with yet not expressed but inherited causal genetic variants, on the accuracy of genetic measures

Age has been commonly used as a covariate in numerous case-control genome-wide association studies. Nevertheless, it is important to note that covariates, while not being influnced by the phenotype of interest (i.e., the outcomes), should function as preventive or promotive factors that exert independent effects on the outcomes [54–56]. In the absence of a standalone impact of age or other variables on the outcome of interest, including these variables as covariates is therefore unwarranted, as it may unnecessarily reduce statistical power [54,55]. In conditions like age-independent disease incidence and in cohorts with wide age ranges, impurity among control samples, i.e., individuals with yet not expressed causal genetic variants, is expected. Therefore, implementing an age cutoff for filtering of control samples, e.g., excluding individuals under 65 years of age, can effectively reduce the impurity levels. It is worth noting that including age as a covariate cannot be considered a corrective measure for predisposed genotypes, as it does not eliminate or reduce the bias introduced by them. The level of impurity among controls depends on the disease prevalence, the disease incidence pattern across different age groups, and the age range and distribution of control individuals within cohorts.

Table 1 provides a simulation of age-independent disease incidence across 10-year age intervals. Here, we used the world population statistics and assumed the same age distribution within the cohort, as well as an even, i.e., age-independent, incidence rate of 12.5% of disease prevalence rate for each 10-year age interval. These conditions result in a control group containing the predisposed genotypes at rates up to ≈ 50% of the population disease prevalence (number of predisposed genotypes, i.e., samples, within the control group = 0.5 × DP × N; DP: disease prevalence, N: total number of samples within the control group). As already mentioned, predisposed genotypes harbour causal genetic variants and will later in life develop the case group phenotype. Therefore, reducing the impurity by filtering out younger individuals from the control group is a reasonable strategy. This typically demands a much larger control group than the case group, a condition common in most genetic cohorts. To demonstrate the potential bias introduced by impurity in control groups and the filtering-by-age efficiency of control samples, we simulated case-control data, based on the information from Table 1 and as discussed in Methods and Text-Box S1. Our analyses revealed a downward bias in odds ratio estimates due to impurity in the control group within cohorts (Figure 4). Simulations at four different population disease prevalence rates of 15%, 10%, 5%, and 1%, each at 9 different SNP-phenotype association levels, revealed the inaccuracy of the estimated odds ratios for the associated SNP (Figure 4). The analyses also indicated the effectiveness of age-dependent filtering within cohorts (Figure 4). We recommend applying the introduced age-dependent filtering of control samples even when dealing with SNPs exhibiting lower association levels, although the downward bias caused by predisposed genotypes is attenuated for SNPs with smaller effect-sizes (Figure 4). This is because, in polygenic scenarios, tens to hundreds of independently associated SNPs, each with small effect sizes, contribute cumulatively, resulting in a substantial downward bias in genetic measures such as polygenic risk scores.

**Figure 4.**
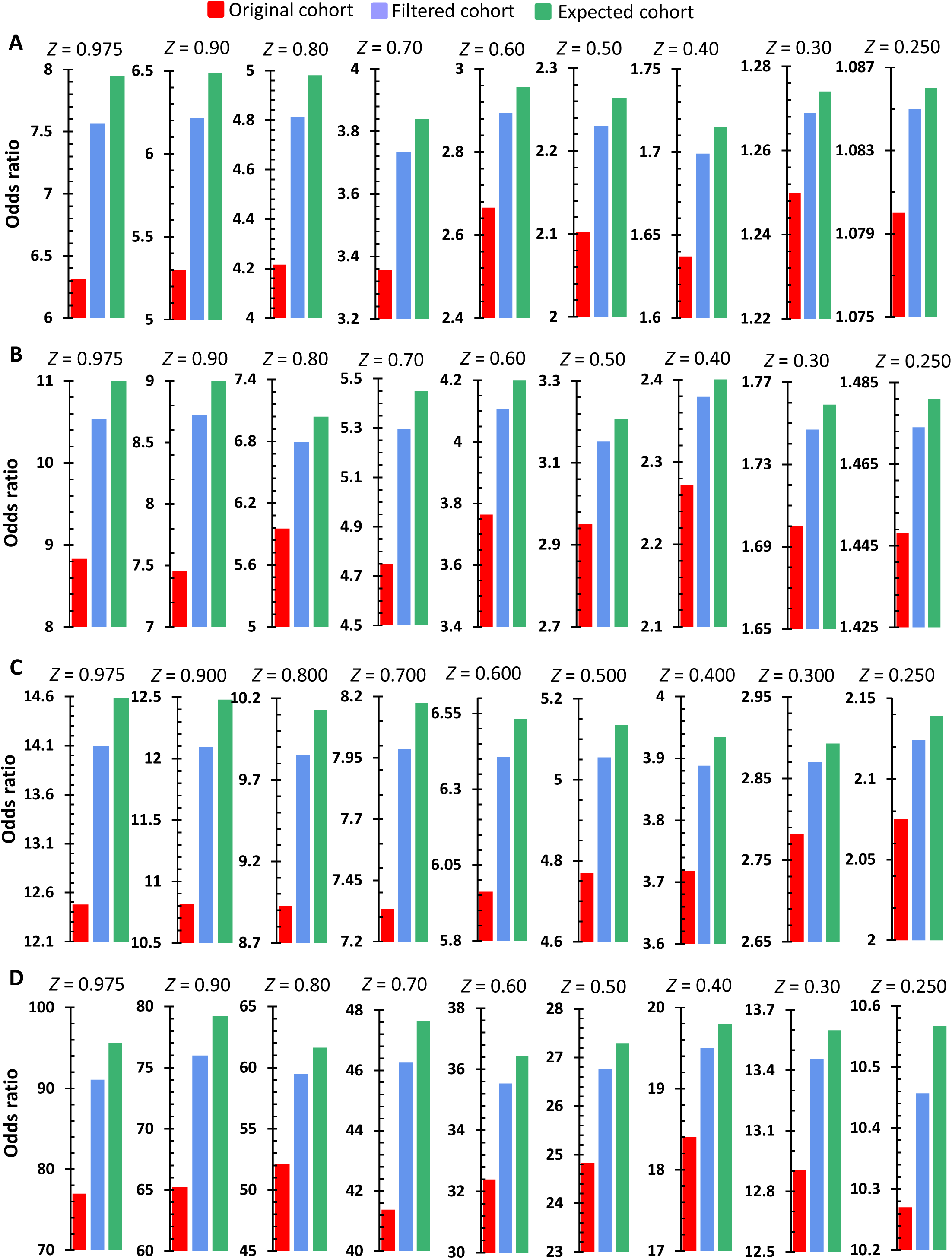
Predisposed genotypes within cohorts introduce downward biases in odds ratios. A) Disease prevalence of 15% and minor allele frequency of 22.5%. B) Disease prevalence of 10% and minor allele frequency of 15%. C) Disease prevalence of 5% and minor allele frequency of 7.5%. D) Disease prevalence of 1% and minor allele frequency of 1.5%. A-D) Genotype simulations were performed as discussed in Methods and Text-Box S1. Based on the global age-distribution and an assumed even incidence rate of 12.5% of disease prevalence for each 10-year age interval, we excluded 90% of controls to simulate filtering of individuals under 65 years of age. Varying levels of association were simulated by applying different levels of Z. For example, Z values of 0.9 and 0.5 indicate that only 90% and 50% of the cases, respectively, are attributable to the causal variant. The control groups within the original cohorts had impurity at half rate of the disease prevalence rates; predisposed individuals were simulated at a half rate of the population disease prevalence as expected and illustrated in Table 1. The control groups within the filtered cohorts had residual impurity at rates of 0.094 × disease-prevalence, as explained in Table 1.

**Table 1.**
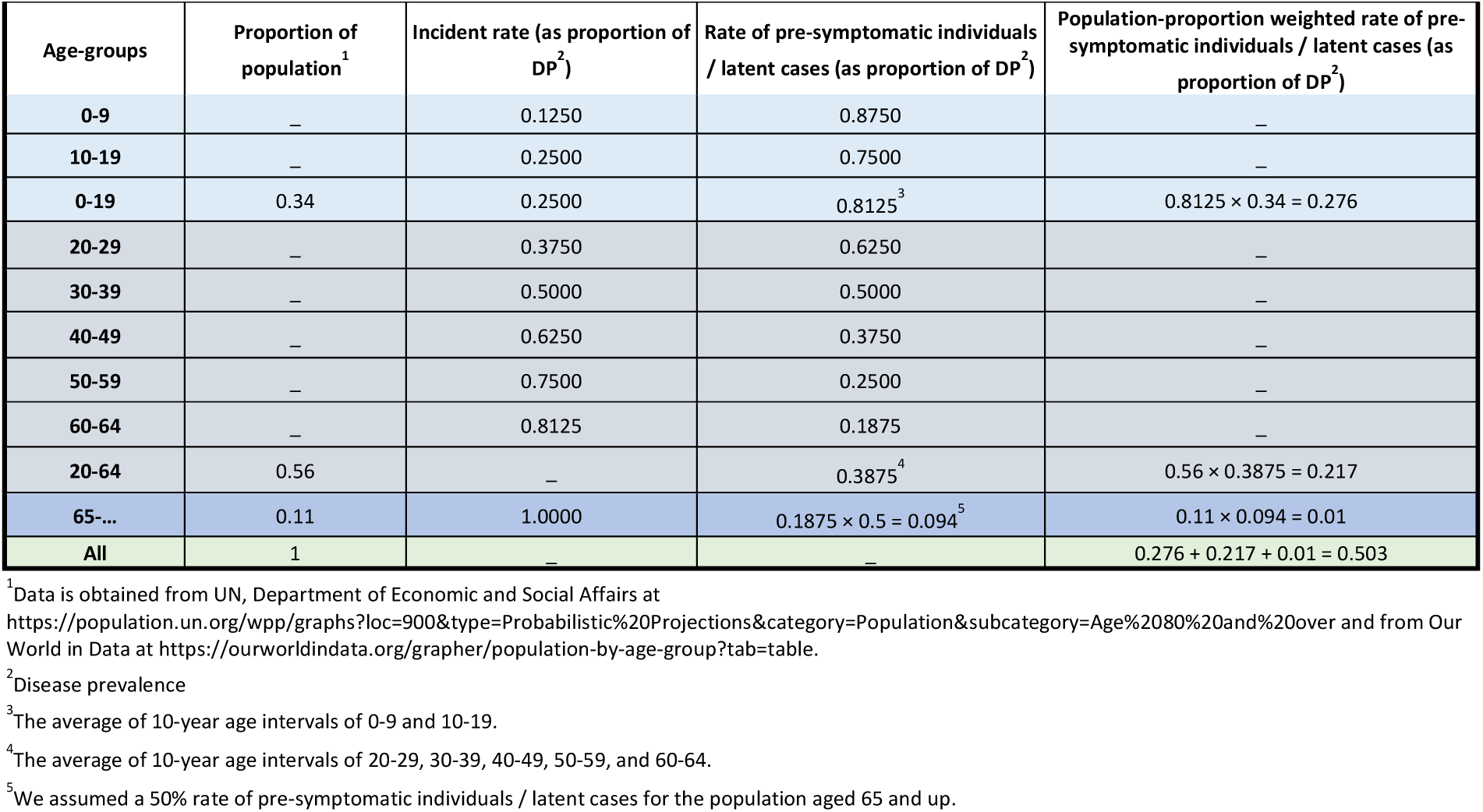
Simulation of predisposed individuals within control groups of cohorts under the assumptions of similar age distribution between the source populations and cohorts and, an even, i.e., age-independent, incidence rate of 12.5% of disease prevalence rate for each 10-year age interval.

It is important to note that the proposed age-filtering method is incompatible with experiments that do not apply the commonly used 1% MAF filter. Cohorts may contain individuals with another specific disease, independent of the study’s disease of interest, which causes early death. Therefore, it is important to highlight that the proposed age-dependent filtering inadvertently depletes the control group of individuals affected by a given disease associated with early mortality. Nonetheless, this should not pose a potential risk of false associations, as a residual number of individuals with an independent second disease linked to early death would still be present among the cases. For example, if the independent second disease, which is entirely associated with early death but excluded from the control group due to age-based filtering, has a prevalence of less than 2%, then the likelihood that cases experienced early death due to this independent disease would also be less than 2%. It should be noted that a 2% prevalence for a disease that always leads to early death is considered exceptionally high and overestimates real-world scenarios. Consequently, genomic variants linked to early death exhibit very low allele frequencies within cohorts; a likelihood of less than 2% among cases is equivalent to a minor allele frequency of less than 1% in a cohort with an equal number of cases and controls. Accordingly, early-death-associated genomic variants are expected to be excluded by applying a quality filter of 1% minor allele frequency.

Furthermore, the death rate among individuals under 65 years old accounts for 40% of all human mortality (Data from Our World in Data at https://ourworldindata.org/grapher/annual-deaths-by-age). Given the same 40% premature mortality rate also among cases, a causal variant would need to contribute to at least 5% of these deaths to have a minor allele frequency of larger than 1% in a cohort with an equal number of cases and controls, following age-related filtering of the control group (0.05 × 0.40 ÷ 2 = 0.01, the division by 2 accounts for the rate being halved after combining the control and case groups). This is also an exceptionally rare condition and overestimates real-world scenarios. Among chronic diseases, only ischaemic heart disease and stroke have global mortality rates exceeding 5%, i.e., account for more than 5% of deaths, including both premature and old-age deaths [6,57,58]. However, these diseases are also highly polygenic, with hundreds of genetic variants being responsible for less than 50% of disease events [6,57]. Taken together, false positive hits originating from the causal genetic variants of early death are not expected after age-related filtering of control groups. This is because a minimum 5% contribution to total deaths in a monogenic scenario and a significantly higher contribution rate in an oligogenic scenario are required for the death-associated genetic variants to pass the minimum 1% minor allele frequency filter. For a polygenic disease that causes early death through hundreds of genetic variants, it is almost impossible to find a causal genetic variant that passes the minimum 1% minor allele frequency filter after age-related filtering of the control group.

It is worth noting that the death rate among individuals under 65 years old in developed countries is significantly lower than the global premature death rate of 40%. This rate for countries like Denmark, Finland, Germany, Italy, Japan, Spain, Sweden, Switzerland, and United Kingdom is below 15% (Data from Our World in Data at https://ourworldindata.org/grapher/annual-deaths-by-age and from Statistics Denmark at https://www.dst.dk/en/Statistik/emner/borgere/befolkning/doedsfald). This underscores the advantage of our proposed age-related filtering: the lower the death rate among individuals under 65, the greater the likelihood that the minor allele frequency filter will exclude variants associated with early death. With 15% of the total deaths occurring at age 0−64, even the extreme assumption that the polygenic ischaemic heart disease which is responsible for 13% of the world’s total deaths [58], is monogenic and that all its related death occur before age 65, cannot lead to false positive hits due to the exclusion of the associated variant after applying 1% minor allele frequency filter (0.15 × 0.13 ÷ 2 = 0.00975 < 1% MAF).

### Non-additive interactions between reference and alternative alleles are a potential source of bias when genetic measures are estimated using the additive-only modeling assumption

Most genome-wide association studies have focused on additive allelic effects, largely because of the prevailing assumption that non-additive effects are negligible [50]. It is important to note that capturing additive-only effects in the presence of both additive and non-additive allelic effects can introduce varying degrees of bias, potentially resulting in false SNP hits and inaccurate genetic measures. We simulated genome-wide association studies (see Text-Box S1 and section Methods) at five different association levels, not only for the additive allelic effect scenario but also for dominance, overdominance, and heterosis-like non-additive allelic effects as described previously [52]. In the presence of non-additive allelic effects, restricting analysis to additive effects led to inaccurate risk scores for genotypes carrying the causal allele, as observed across five association levels and two disease prevalence rates (Table 2). However, this is not the case when applying the allelic adjustment method, which estimates the true allelic effect using the lowest association *p*-value for each SNP across different effect-model scenarios [52].

**Table 2.**
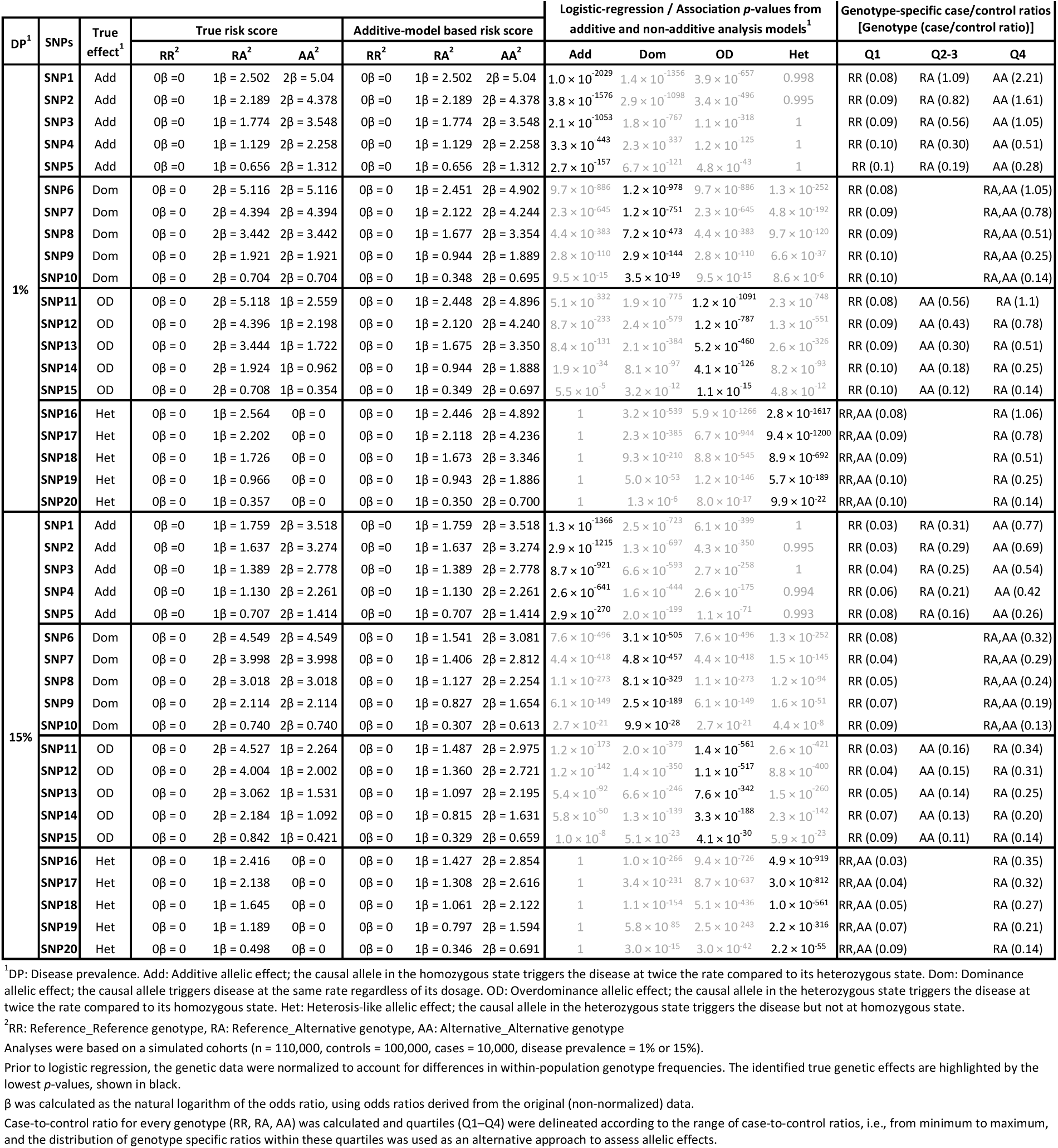
Additive-model based association analyses tend to produce inaccurate risk score estimates under conditions of non-additive allelic interactions.

To jointly capture additive and non-additive allelic effects, we applied the SAFE-*h*^2^ allelic adjustment approach, which uses observed allelic effects and subsequent allelic adjustments to fit the observed allelic effect-models [52]. This means that every SNP is analyzed according to its specific allelic interaction model, i.e., whether additive or one of the non-additive models. The advantage of the mentioned allelic adjustment approach is its reliance on a linear association model while effectively capturing both additive and non-additive allelic effects. Here, it is important to highlight the methodological difference between quantitative and qualitative traits. For quantitative traits, it is the quantities, including every observation, that influence the regression output. In contrast, the frequency of each phenotypic category plays an important role in the logistic regression output. Logistic regression, which is used for qualitative traits, is therefore sensitive to the frequency of each phenotypic category relative to the genotypes, i.e., the three SNP-specific genotypes (Ref/Ref, Ref/Alt, and Alt/Alt) observed for both the control and case groups. Due to differences in genotype (Ref/Ref, Ref/Alt, and Alt/Alt) frequencies within the population, the sensitivity to the frequency of phenotypic categories hinders the identification of the best-fitted model when comparing allelic interaction models. Therefore, we introduce a normalization step which is essential when analyzing qualitative phenotypes to account for differences in genotype frequencies within the population. This process ensures a fair comparison of allelic interaction models, enabling the identification of the model that most accurately represents the observed data and provides the best fit. For normalization, genotype frequencies within the population are required, as explained in section Methods and Text-Box S2. Assuming known population allele frequencies, we can calculate population genotype frequencies (see Text-Box S1) and use these values to normalize the observed genotype frequencies (see Methods and Text-Box S2). If within-population allele frequencies are unknown, the population disease prevalence can be used to build iterative control-case combinations, each representing the original population. Thus, the average genotype frequencies can be used as estimates of population genotype frequencies. Please note that normalization is relevant only when comparing allelic interaction models to identify the best model for each SNP. The original unnormalized data should be used, as previously described, for the allelic adjustment strategy to obtain genetic measures for SNPs with additive allelic effects and for SNPs exhibiting non-additive allelic effects.

In contrast to the common approach, i.e., additive-effects based association analysis, which performs poorly in the presence of non-additive allelic interactions and yields inaccurate risk scores, the SAFE-*h*^2^ allelic adjustment approach enabled clear differentiation of the allelic interaction models (Table 2). Here, we simulated and applied predesignated true associations using various genetic effect models. By evaluating different effect-model scenarios for every SNP, the lowest association *p*-values were obtained in the scenario that matched the true allelic effect (Table 2). In Table 2, the expected (true) risk scores also corresponded to those obtained using the allelic adjustment method. The results indicate that the allelic adjustment strategy, when combined with normalization can be applied in case-control studies to achieve greater accuracy. SAFE-*h*^2^ performs model selection based on association *p*-values, effectively distinguishing among additive and various non-additive genetic effects at varying association levels. In contrast to association *p*-values (Table S1), the Akaike Information Criterion (AIC) and Bayesian Information Criterion (BIC) failed to distinguish between predesignated genetic models of additive and overdominance (Table S2).

The normalization of genetic data to correct for differences in population genotype frequencies and the subsequent logistic regression for every SNP are considerable tasks. A more practical and efficient approximation is to leverage the case-to-control ratios for each genotype count (Ref/Ref, Ref/Alt, & Alt/Alt). Afterwards, we can define quartiles (Q1 to Q4) by dividing the range of case-to-control ratios into four equal parts. Finally, the distribution of genotypes across these quartiles can be used to define allelic effects as shown in Table 2. We found complete agreement between allelic effects identified by association *p*-values obtained after normalization of genotype frequencies and those inferred from case-to-control ratios of genotypes (Table 2 and Table S3). To facilate the identification of allelic effects using case-to-control ratios of genotypes, we developed the Allelic-effect aware Case-Control GWAS tool (AlleliC-GWAS). AlleliC-GWAS is an open-source tool that reads plink binary files as input, where the MAIN.fam file contains binary traits encoded as 0 vs 1 or 1 vs 2, to identify SNP-specific allelic effects. Allelic effects include additive, tagged as [Add], dominance effects by Ref and Alt alleles, tagged as [Dom] and [dom], overdominance effects by Ref and alternative alleles, tagged as [OD] and [od], and heterosis-like effects, tagged as [Het]. AlleliC-GWAS output also includes a combinatory bfile with adjusted allelic effects, along with association statistics generated using the PLINK GLM function for the adjusted SNPs. Users can also use the generated bfiles for other GWAS analyses, including GCTA based mixed model [23] and conditional association tests [59] as well as the SAFE-*h*^2^ based SNP heritability estimation [52].

### Accurate analysis of multi-allelic genomic loci necessitates decomposing variants into individual allelic comparisons relative to a defined reference allele

In comparison to biallelic SNPs, there is a higher complexity in the association analyses of multi-allelic SNPs. Higher levels of genotypic diversity cause this complexity, which can be further exacerbated by non-additive allelic effects or by a combination of additive and non-additive allelic effects. Given the same cohort size, greater genotypic diversity at multi-allelic sites also reduces statistical power relative to biallelic SNPs in association analyses. Simultaneous association analysis for all alleles of a multi-allelic SNP demands complex models. The most common coding format for multi-allelic SNPs only allows comparison of each allele versus the remaining alleles collectively as the reference. This particular allele coding format may introduce an additional source of bias. Comparing every allele versus the remaining alleles of a multi-allelic genomic position can elevate false negative rates and compromise the accuracy of estimates for the allele’s genetic effects. The expected inaccuracy is due to possible contrasting genetic effects among the alleles, collectively defined as the reference allele. Therefore, an ideal solution is to set one of the alleles, e.g., the most frequent allele, as the reference allele and perform association analyses for the remaining alleles, each versus the reference allele. In addition to the association analyses using a collective reference allele for multi-allelic genomic positions, we accordingly recommend performing an association analysis for each allele in a multi-allelic genomic position versus a defined single-allele reference. For example, four association analyses are performed for a genomic position with four alleles. In all these four association analyses, the reference allele is a combination of three alleles. In this example, we recommend conducting three additional association analyses, each using the most frequent allele as the reference, and comparing it individually against each of the other three alleles. This is feasible through reformatting the dataset to decompose multi-allelic genomic positions.

## Discussion

A key challenge in genome-wide association studies is inadequate genotypic coverage; approximately 3^N^ individuals with distinct genotypes are required for precise genetic estimation in a diploid genome with N biallelic SNPs, given that mutations are independent and conditional lethal SNPs are exceedingly rare. Meeting such cohort sizes is unfeasible due to the relatively small size of the human population. For instance, insufficient genotypic coverage is the reason for regional association signals from multiple neighbouring SNPs, and the developed SNP clumping and fine-mapping strategies [60–65] only partially overcome this challenge by identifying the most likely causal SNP or SNP_s_ within each regional association signal. Replicating experiments, e.g., using effect-sizes of associated SNPs obtained from one cohort to calculate polygenic risk scores for individuals in a second cohort, is also a common approach to mitigate overfitting caused by inadequate genotypic coverage. Other factors may also introduce bias into the experiments, e.g., the misclassification of predisposed genotypes as control samples, genetically related samples within cohorts [13–20], and non-additive allelic interaction [52]. Unlike the challenge posed by insufficient genotypic coverage, which is an insurmountable obstacle, the remaining factors can be efficiently addressed to reduce bias in genetic assessments.

Our results indicate the disadvantage of the standard bulk filtering procedure for genetically related individuals within cohorts. We present evidence that genetic relatedness across case-control groups effectively reduces false positive rates and, therefore, should not be removed through bulk filtering. Ideally, the only source of variation among experimental groups or treatments should be the phenotype of interest. We often include covariates to account for known sources of variation, allowing for a less biased comparison of samples with respect to the phenotype of interest. Genetic relatedness across case-control groups serves a similar purpose by controlling for both known and hidden sources of variation, thereby establishing an experimental framework for more robust and unbiased evaluation of the phenotype of interest among samples. In other words, genetic relatedness between case and control groups selectively reduces allele-frequency differences for non-causal variants, where all observed disparities are attributable to experimental bias. Considering 25% genetic relatedness, we found no bias leading to false-positive hits attributable to genetically related sub-communities constituting up to 0.25% of controls or cases, or by genetically related paired-samples constituting less than 10% of the case or control groups. The mentioned thresholds of 0.25% and 10% impose a high degree of stringency, given that the simulated 25% genetic similarity exceeds typical expectations and represents the upper-bound of relatedness levels in real-world cohorts.

The proposed age-dependent filtering also effectively mitigated bias caused by predisposed genotypes that were incorrectly classified as control samples. Age-based filtering is applicable when age is not a covariate and, as a characteristic commonly observed in contemporary genetic cohorts, the control group is substantially larger than the case group. This method can thus be effectively employed to correct for bias arising from predisposed genotypes. In general, age-based filtering is incompatible with experiments that omit the commonly used 1% MAF filter, as this may lead to false positives, particularly for phenotypes with high within-population prevalence. Nevertheless, age-based filtering can still be applied in rare variant analyses by conducting a complementary GWAS using age-based filtered samples, i.e., deceased samples versus the remaining filtered samples. This complementary GWAS can identify potential variants linked to premature death that might otherwise yield false positives after age-based filtering in rare variant analyses.

Cox regression is a robust method for analyzing time-constant variables that influence time-to-event outcomes [66,67]. When disease age-of-onset is defined as a time-to-event phenotype to perform survival analysis in GWAS, individuals/genotypes are treated as components of a time-constant variable. Here, special attention should be given to the age of onset, as it requires a strong correlation between disease incidence and specific ages. This correlation should also result from a direct effect of age on the outcome, i.e., disease incidence. In other words, defining age-of-onset as a time-to-event phenotype requires that age serve as a covariate that mediates the genetic influence on disease incidence. Otherwise, disease incidence across all ages, i.e., with no clear correlation or enrichment at specific ages, reflects the dependence of disease age-of-onset on non-genetic factors. For example, we have found an even distribution of incidence for inflammatory bowel disease across all ages [53]. Notably, Cox regression cannot account for predisposed genotypes, i.e., impurity in control groups, where disease incidence is driven by non-genetic factors. Predisposed genotypes consistently lead to underestimation of effect-sizes; consequently, accurate inference methods should adjust for this bias and report appropriately larger effect-sizes. However, Cox regression often leads to smaller effect-sizes. For instance, when conducting association analyses for coronary heart disease, Cox regression yielded smaller effect-size estimates compared to logistic regression [66]. In agreement, the hazard ratios from Cox regression on the simulated data (see figure 4 for the dataset) were also smaller than the odds ratios (Figure S3).

Genome-wide association studies have predominantly modeled additive allelic effects, reflecting the commonly held assumption that non-additive effects contribute minimally to trait variations [50]. However, there is no inherent genetic or biological rationale to rule out non-additive allelic effects. Given the roles of evolutionary processes through random genetic mutations and the complexity of molecular interactions at all levels that modulate biosynthetic, structural, and compositional properties, the occurrence of non-additive allelic effects is highly plausible. Here, as an example, we refer to two recent studies on dominance effects in mammals [47] and humans [48]. Methodological limitations have been introduced as the main driver for the trivial estimates of non-additive allelic effects [51,52]. We further introduce the impact of population-level frequency disparities among SNP genotypes (Ref/Ref, Ref/Alt, Alt/Alt), which complicate the identification of allelic interaction modes for individual SNPs via logistic regression in case-control analyses. The results indicate that a pre-normalization step for SNP-specific genotype frequencies enables accurate detection of allelic interaction modes and thereby reduces bias in capturing phenotypic variance. Here, we introduce AlleliC-GWAS tool to identify allelic effects on binary traits using case-to-control ratios of genotypes. It is important to note that capturing additive-only effects in the presence of additive and non-additive allelic effects can introduce varying degrees of bias, potentially resulting in false-negative hits and inaccurate effect-size and other genetic measures.

## Conclusions

Predisposed genotypes lead to downward bias in effect-size estimates. The bias introduced by predisposed genotypes can be minimized by excluding young individuals from control cohorts. Bulk filtering of genetically related samples is not an ideal approach; instead, filtering related samples found within case or control groups while retaining genetically related events shared between case and control groups helps reduce false-positive rates. Furthermore, the lack of two-allele comparisions at multi-allelic geomic positions can potentially lead to false-negaive results in association tests, highlighting the importance of proper allele decoding. Finally, the AlleliC-GWAS tool is designed to provide accurate genetic measures for binary traits by jointly capturing additive and non-additive allelic effects after normalization of SNP-specific genotype frequencies.

## Materials and methods

### Simulation of data

We used LDAK to simulate unrelated genotypes [68]. Afterwards, unrelated individuals were applied to simulate genetically related individuals. We replicated ≈ 25% of the SNPs from one individual in an unrelated individual to obtain two genetically related individuals. As expected, genetic similarity among related individuals was 24.8%–25.1%, calculated using the KING-robust kinship estimator of PLINK [14]. To analyze the relatedness-based filtering, case and control allocations were randomized across individuals. Therefore, any increase in the number of significant SNP hits, e.g., due to the inclusion of genetically related case or controls, indicates an elevated false-positive rate. On the other hand, any decrease in the number of significant SNP hits, e.g., due to the inclusion of individuals that introduce genetic relatedness across case and control groups, indicates an reduced false-positive rate. However, any decrease in the number of significant SNP hits in the absence of an underlying cause, i.e., without any related samples being shared between the case and control groups, is an indication for increased false negative rates. We performed the analyses in 60 independent replicates. Association analyses were performed using the GLM function of PLINK [69]. Minor allele frequencies were ≥ 0.01 in all the analyses. For mixed model based analysis, GCTA-fastGWA-GLMM was used [23].

The age distribution of the simulated cohorts was based on the world age distribution, as shown in Table 1. Varying levels of true association were simulated by applying a factor, *Z*, representing the percentage of disease prevalence attributable to the causal variant. Disease prevalence rates were simulated at 15%, 10%, 5%, and 1%, with corresponding causal allele frequencies of 22.5%, 15%, 7.55%, and 1.5%, respectively. The simulations required causal allele frequencies to exceed disease prevalence to account for predisposed genotypes, i.e., individuals carrying the causal alleles but not yet affected, within the control group. The calculation of population and cohort level genotype frequencies for genotypes within population (XX_P_), genotypes of healthy controls after excluding cases from the population (XX_PH_), case genotypes within the cohort (XX_C_), control genotypes within the cohort (XX_H_), and control genotypes within the cohort after removing individuals under 65 years of age (XX_H_) were performed as follows, in steps 1 to 5:

1. *Population* (*Disease prevalence = 1%, Frequency of Reference allele A = 0.985 & frequency of causal allele C = 0.015*) Genotype frequencies →

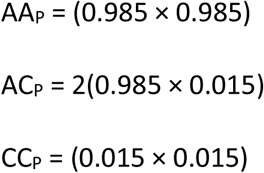 Patient and non-patient frequencies → Healthy_P_ = 0.99, Healthy with predisposed genotypes = 0.005 Patients_P_ = 0.01
2. *Calculating genotype (i.e., AA, AC, & CC) frequencies for cases witihin cohort* (*n = 10,000: 0.0% controls + 100% cases, proportion of cases being associated with allele C = Z*) Causal-allele C independent cases →

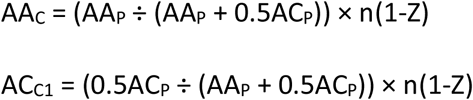 Causal-allele C dependent cases →

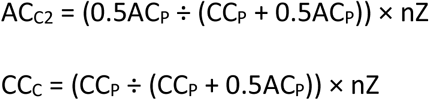
3. *Calculating genotype, i.e., AA, AC, & CC, frequencies for pure (with no predisposed genotype) controls* (*n = 100,000: 100% controls + 0.0% cases*)

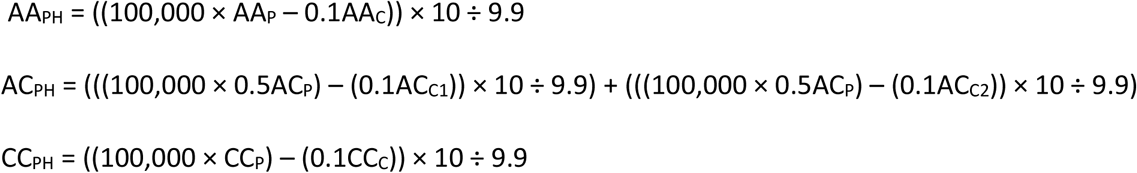
4. *Calculating genotype, i.e., AA, AC, & CC, frequencies for controls within cohort* (*n = 100,000: 99.5% controls + 0.5% cases*)

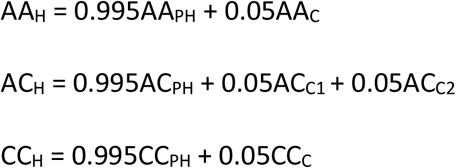
5. *Calculating genotype, i.e., AA, AC, & CC, frequencies for filtered controls* (*age ≥ 65 years*) *within cohort* (*n = 10,000: 99.99% controls and 0.01% cases*)

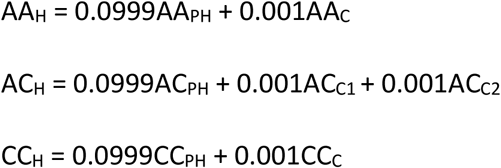

The same procedure was applied for simulations with disease prevalences of 5%, 10% and 15%.

### Additive and non-additive allelic interactions

Here, we adopted the approach described above to simulate and apply predesignated true associations, this time using various genetic-effect models. We used SAFE-*h*^2^ for joint-capturing of additive and non-additive allelic effects [52]. This is done through allelic adjustments for linear approximation of allelic effects, including dominance for either the reference or alternative allele, overdominance for either the reference or alternative allele, and heterosis-like effects. When linear models are augmented with some allelic adjustments tailored to the observed non-additive effects, e.g., by converting the heterozygote nucleotide positions into one of the homozygotes at the genomic positions with dominance allelic effects, we can capture genetic variance at levels comparable to those of the exponential model. SAFE-*h*2 employs the same allelic adjustment algorithm to extract non-additive effects within the framework of linear models as described previously [52]. Briefly, SAFE-*h*^2^ examines all scenarios, i.e., additive (original genotypes), adjusted genotypes for dominance effects, adjusted genotypes for overdominance effects, and adjusted genotypes for heterosis-like effects, to find the best-fitted linear model for every SNP position. Before association analysis, combinatory genotype datasets can be built based on a selected adjustment or the original genotype, (i.e., the one with the smallest association *p*-value, for each SNP position). In case-control studies employing logistic regression for genome-wide association analyses, the SAFE-*h*2 allelic adjustment algorithm requires a pre-normalisation step. In contrast to the linear regression on quantitative phenotypes, where the quantity of every genotype is highly influential, logistic regression is sensitive to the frequency of each phenotypic category in relation to the genotypes, i.e., SNP specific three genotypes of Ref/Ref, Ref/Alt, Alt/Alt among control samples and among case samples. Therefore, population level frequency differences among SNP specific genotypes of Ref/Ref, Ref/Alt, and Alt/Alt can hinder the identification of the best fitted model when comparing allelic interaction models through logistic regression. Therefore, a normalization step is necessary when analyzing qualitative phenotypes to account for genotype frequency differences within population before being able to compare the allelic interaction models and identify either additive or one of the non-additive models that provides the best fit. The correction for genotype frequency differences can be carried out as outlined below:

Considering XX_P_ for genotype frequencies within population, XX_C_ for case-samples specific genotype frequencies within the cohort, XX_CN_ for normalized case-samples specific genotype frequencies within the cohort, XX_H_ for control-samples specific genotype frequencies within the cohort, and XX_HN_ for normalized control-samples specific genotype frequencies within the cohort, we correct the frequencies as follows, in steps 1 and 2:

1. *Population* (*Reference Allele frequency = R, Alternative Allele frequency = A*)

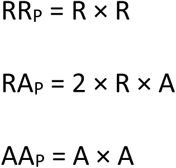

When within-population allele frequencies are unknown, the population disease prevalence can be used for iterative control-case combinations, each representing the original population. In this way, the average genotype frequencies will provide estimates for population genotype frequencies.
2. *Cohort* *2-1.* Observed genotype frequencies among controls within the cohort:

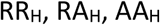 Normalized genotype frequencies among controls within the cohort (Assuming R > A):

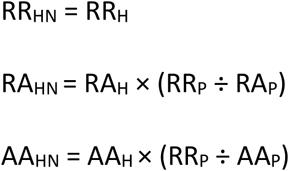
3. *2-2.* Observed genotype frequencies among cases within the cohort:

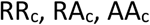 Normalized genotype frequencies among cases within the cohort (Assuming R > A):

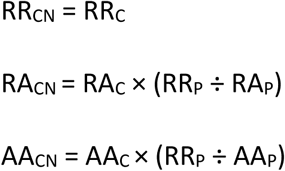

This normalization creates a new cohort dataset that can be used to determine the best-fitting interaction models for alleles of each SNP using the SAFE-*h*^2^ allelic adjustment algorithm. Finally, the identified interaction models for each SNP can be applied to the original dataset (not normalized) using the SAFE-*h*^2^ allelic adjustment algorithm to obtain accurate genetic measures.

As an efficient approximation of the regression line, we also used the case-to-control ratios (CCR) for SNP-specific genotypes (CCR_Ref/Ref_, CCR_Ref/Alt_, CCR_Alt/Alt_). In rare instances of complete association or when predisposed genotypes are absent among control samples, certain genotypes may be missing from either the control or case group. To address this and avoid zero or undefined ratios, a small numeric constant, e.g., 10^-6^, should be introduced before obtaining ratios. Four quartiles were defined for the whole range of CCR, i.e., Max_CCR − Min_CCR. Finally, the distribution of SNP-specific CCR_Ref/Ref_, CCR_Ref/Alt_, and CCR_Alt/Alt_ ratios across these quartiles was used to define allelic effects as shown in Table 2. The additive model can be confirmed when CCR_Ref/Ref_, CCR_Ref/Alt_, and CCR_Alt/Alt_ are distributed respectively in Q1, Q2-Q3, and Q4, or alternatively in Q4, Q2-Q3, and Q1. The dominance model can also be confirmed when CCR_Ref/Ref_ falls within Q1, while both CCR_Ref/Alt_ and CCR_Alt/Alt_ fall within Q4, or when CCR_Ref/Ref_ falls within Q4, while both CCR_Ref/Alt_ and CCR_Alt/Alt_ fall within Q1. Alternatively, the dominance model can be supported when CCR_Ref/Ref_ and CCR_Ref/Alt_ fall within Q1 while CCR_Alt/Alt_ falls within Q4, or when CCR_Ref/Ref_ and CCR_Ref/Alt_ fall within Q4 while CCR_Alt/Alt_ falls within Q1. The overdominance model is confirmed when CCR_Ref/Ref_, CCR_Alt/Alt_, and CCR_Ref/Alt_ are distributed respectively in Q1, Q2-Q3, and Q4; or in Q2-Q3, and Q4, and Q1; or alternatively in Q4, Q2-Q3, and Q1; or in Q2-Q3, Q1, and Q4. The heterosis-like model can also be supported when CCR_Ref/Ref_ and CCR_Alt/Alt_ fall in Q1 while CCR_Ref/Alt_ in Q4, or alternatively when CCR_Ref/Ref_ and CCR_Alt/Alt_ fall in Q4 and CCR_Ref/Alt_ falls in Q1. We further developed the open-source tool “AlleliC-GWAS” to perform the normalization for genotype frequencies by applying genotype count case-to-control ratios. AlleliC-GWAS requires plink binary files as input. The “fam” file can contain binary traits encoded as 0 vs 1 or 1 vs 2. AlleliC-GWAS identifies allelic effects of additive [Add], dominance effects by Ref allele [Dom] and Alt allele [dom], overdominance effects by Ref allele [OD] and alternative [od], and heterosis-like effects [Het]. AlleliC-GWAS provides a combinatory bfile with adjusted allelic effects, along with association statistics on the adjusted SNPs.

## Contributions

BD and VA conceived the study. BD, OBVP, SRO, QT, and VA discussed the experimental designs and results. BD performed the analyses. BD and VA wrote the manuscript in consultation with OBVP, SRO, and QT.

## Acknowledgements

Behrooz Darbani acknowledges Louis-Hansens Fond (grant nr. 23-2B-14148). Vibeke Andersen acknowledges Sundhedsdonationer (grant nr. 2024-0379) and Region of Southern Denmark (grant nr. 23/7359). The authors thank Caroline Moos (Research advisor, Department of Clinical research, Hospital Sønderjylland) for invaluable language-editing contribution.

## Availability of data and materials

The datasets supporting the conclusions of this article are included within the article and its additional file. AlleliC-GWAS tool and its associated manual is available for download at https://github.com/SAFE-h2/AlleliC-GWAS.

## Supplementary information

**Figure S1.**
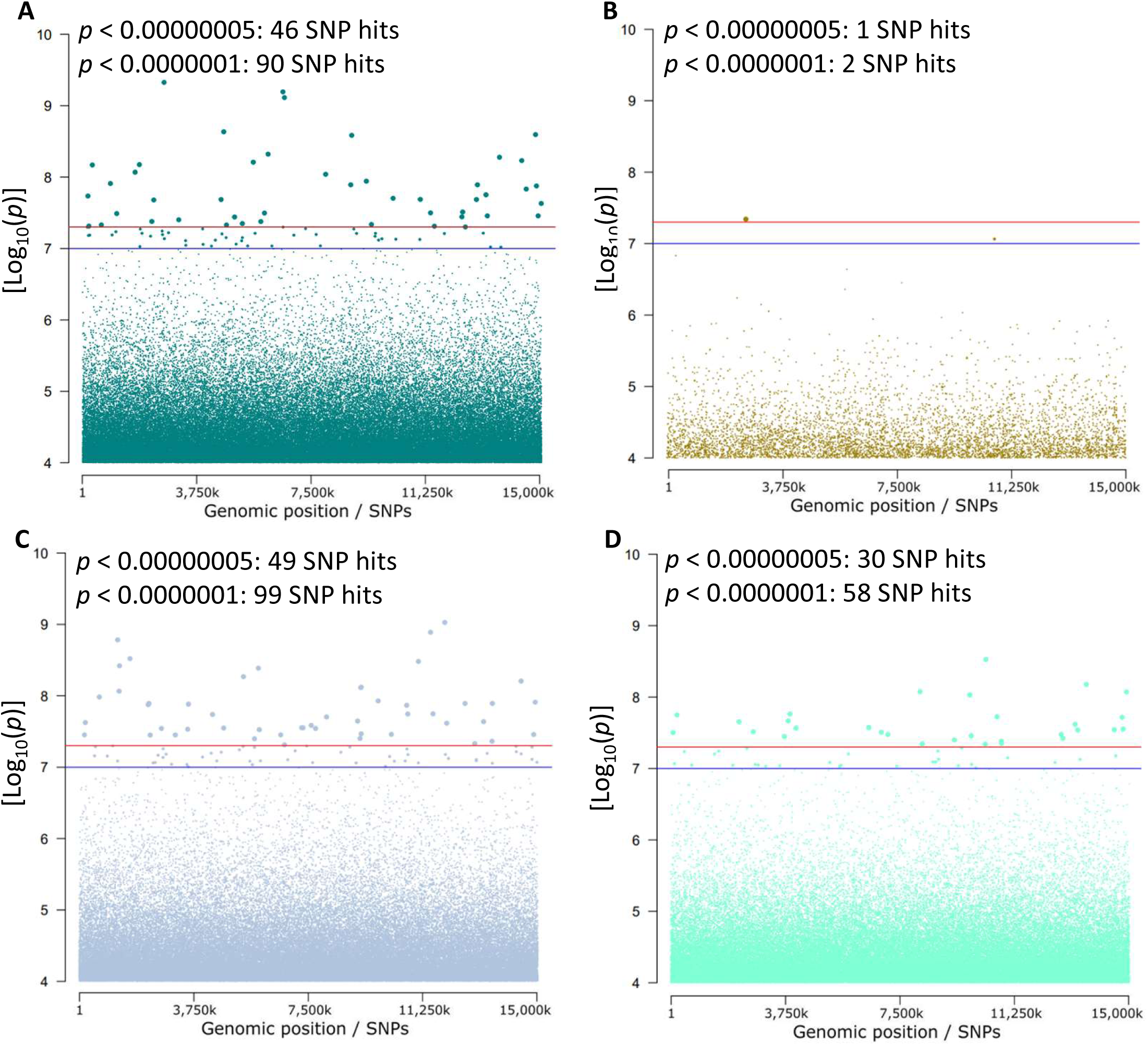
Random associations across varying numbers of genetically related, case- or control-group specific paired samples as well as relatedness between case and control groups. The generalized linear mixed model of fastGWA-GLMM was applied for association analyses. Each Manhattan plot is illustrating the outputs from 60 independent replicates of case-control association analyses. The analyses were based on simulated data including 15 million SNPs (MAF ≥ 0.01) and 20,000 samples. Random association signals in these datasets reflect false positive rates resulting from random allocation of case and control samples. Moreover, a smaller number of random associations observed in scenarios with case- or control-specific relatedness (D)—compared to scenarios without such relatedness and also without relatedness shared across case and control groups (A)—indicates an increased risk of false negatives. A) Analyzing genetically unrelated samples. Genetic similarities among cases, controls, and case-control samples were ≤ 0.3%. B) Analyzing genetically related samples. Genetic similarities between case and control samples were 24.8%–25.1%. Genetic similarities among cases and among controls were ≤ 0.3%. C) Analyzing genetically unrelated samples. Genetic similarities among cases, controls, and case-control samples were ≤ 0.3%, except for a sub-community of 50 control samples, each sharing 24.8% to 25.1% genetic similarity with all other samples within the sub-community. D) Analyzing genetically unrelated samples. Genetic similarities among cases, controls, and case-control samples were ≤ 0.3%, except 1,000 paired samples from the control group exhibiting 24.8%–25.1% genetic similarity between paired samples. MAF: Minor allele frequency.

**Figure S2.**
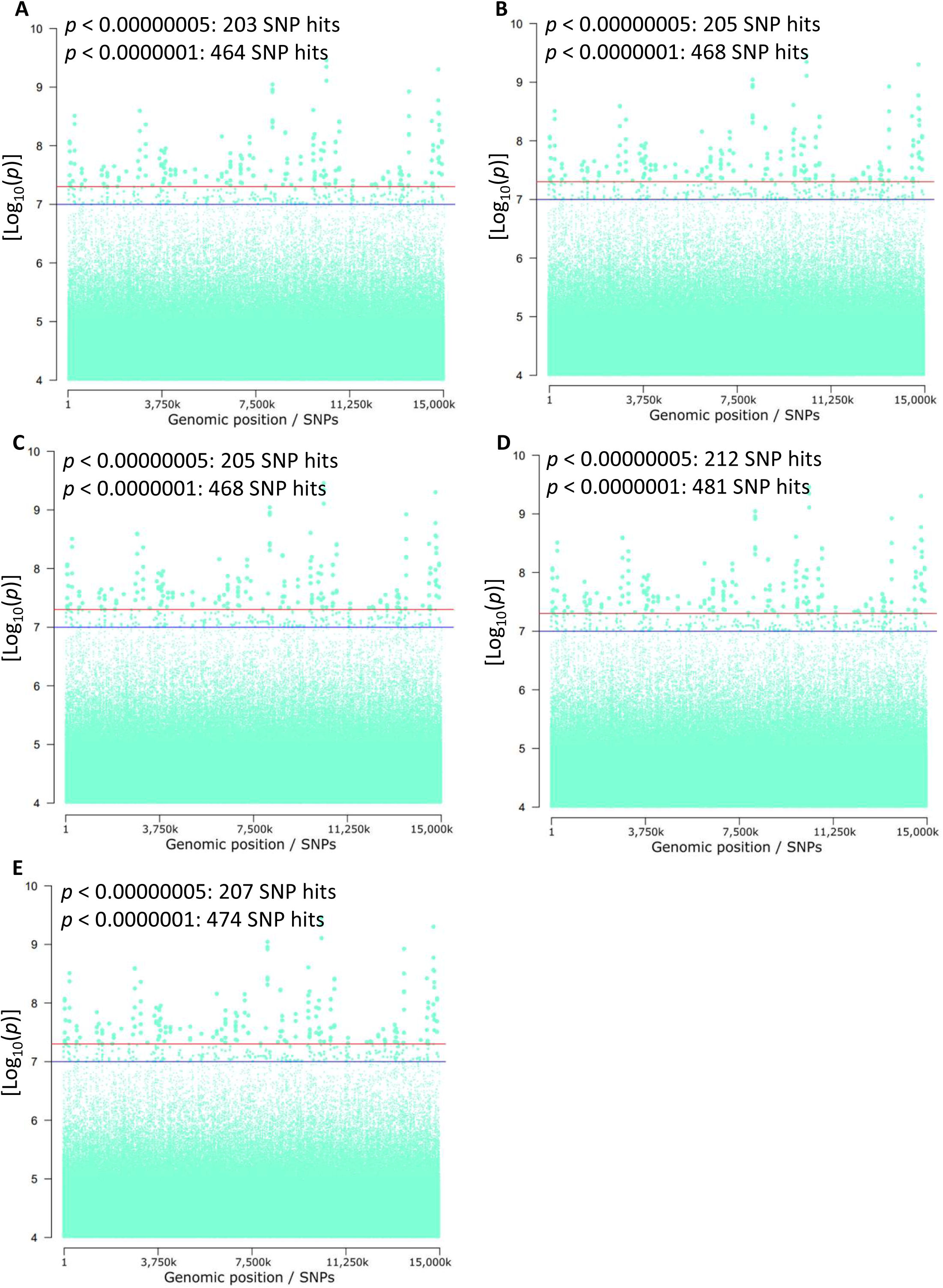
Random associations after filtering control-group specific related samples. Sample filtering was performed five times (A-E), each by excluding 1000 samples from 1000 related paired-samples leaving no relatedness. The generalized linear mixed model of fastGWA-GLMM was applied for association analyses. Each Manhattan plot is illustrating the outputs from 60 independent replicates of case-control association analyses. The analyses were based on simulated data including 15 million SNPs (MAF ≥ 0.01) and 20,000 samples (19,000 samples after sample filtering). Genetic similarities among cases, controls, and case-control samples were ≤ 0.3%, except 1,000 paired samples from the control group exhibiting 24.8%–25.1% genetic similarity between paired samples. MAF: Minor allele frequency.

**Figure S3.**
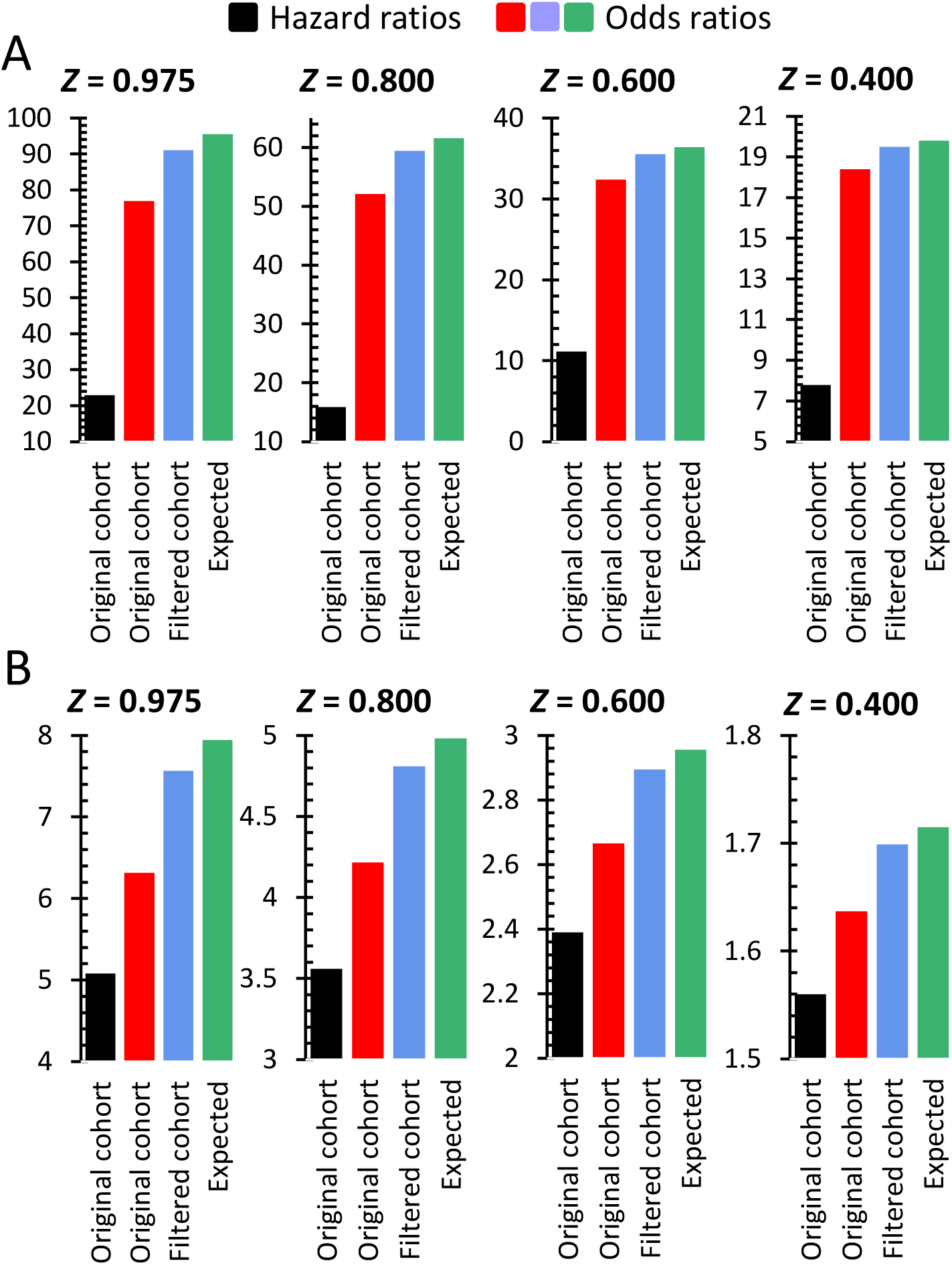
The presence of predisposed individuals within cohorts introduces downward bias in odds ratio estimates. A) Disease prevalence of 1% and minor allele frequency of 1.5%. B) Disease prevalence of 15% and minor allele frequency of 22.5%. A,B) Genotype simulations were performed as discussed in Methods and Text-Box S1. Based on the global average age-distribution and an even incidence rate of 12.5% of disease prevalence for each 10-year age interval, we excluded 90% of controls to simulate filtering of individuals under 65 years of age. Varying levels of association were simulated by applying different levels of Z. For example, Z values of 0.8 and 0.4 indicate that only 80% and 40% of the cases, respectively, are attributable to the causal variant. The control groups within the original cohorts had impurity at half rate of the disease prevalence rates; predisposed individuals were simulated at a half rate of the population disease prevalence as expected and illustrated in Table 1. The control groups within the filtered cohorts had residual impurity at rates of 0.094 × disease-prevalence, as explained in Table 1. Age was used as time-to-outcome in Cox regression.

**Text-Box S1.**
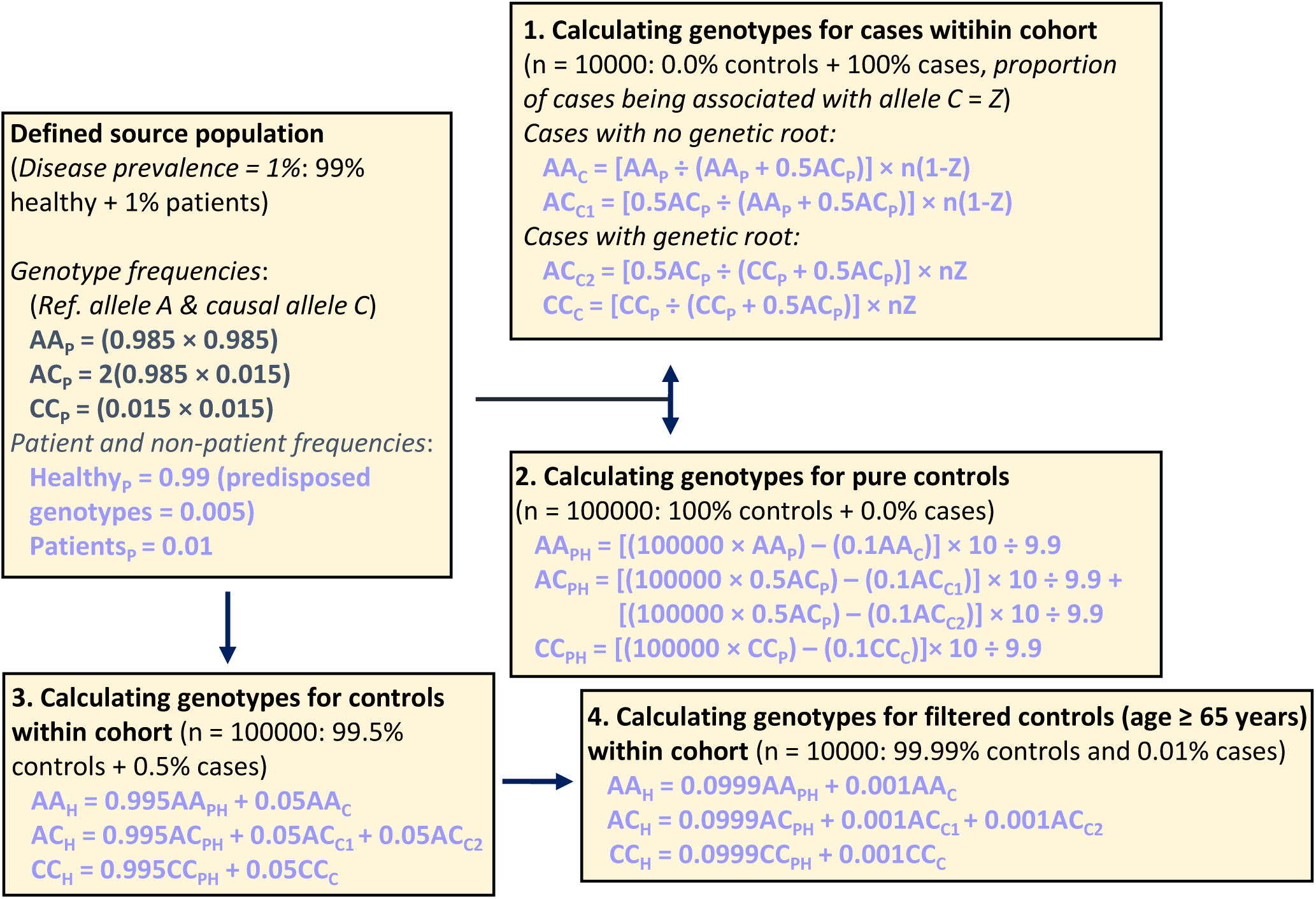
Simulation of genotypes within population (XX_P_), genotypes of healthy controls after excluding cases from the population (XX_PH_), case genotypes within the cohort (XX_C_), control genotypes within the cohort (XX_H_), and control genotypes within the cohort after removing individuals under 65 years of age (XX_H_). According to Table 1, individuals younger than 65 years accounted for 90% of the original cohort. Varying levels of association were simulated by adjusting the value of Z, which can range from 1 (representing the full rate of disease prevalence) to 0 (representing no contribution to disease prevalence). For example, Z values of 0.9 and 0.5 indicate that 90% and 50% of the cases, respectively, are attributable to the causal variant. Here, a simulation at 1% disease prevalence and 1.5% causal allele frequency is shown. Similar simulations were also performed at 5% disease prevalence and 7.5% causal allele frequency, 10% disease prevalence and 15% causal allele frequency as well as at 15% disease prevalence and 22.5% causal allele frequency.

**Text-Box S2.**
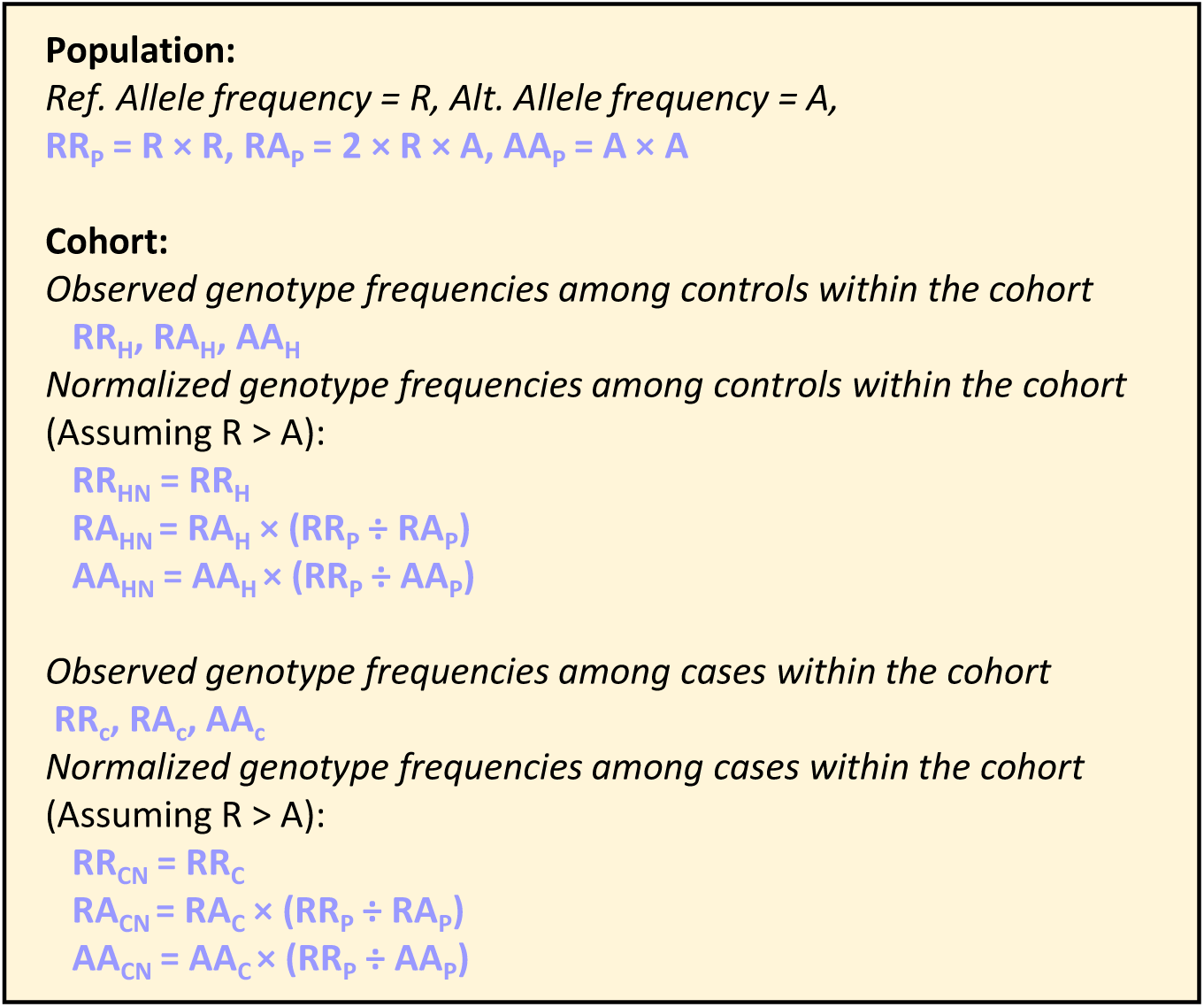
Normalization of genetic data for differences of genotype frequencies within the population prior to examining the allelic interaction models for each SNP. This normalization is required for qualitative traits to be analyzed through logistic regression. Genotype frequencies within population (XX_P_), case-samples specific genotype frequencies within the cohort (XX_C_), normalized casesamples specific genotype frequencies within the cohort (XX_CN_), control-samples specific genotype frequencies within the cohort (XX_H_), and normalized control-samples specific genotype frequencies within the cohort (XX_HN_).

---

**Table S1.**
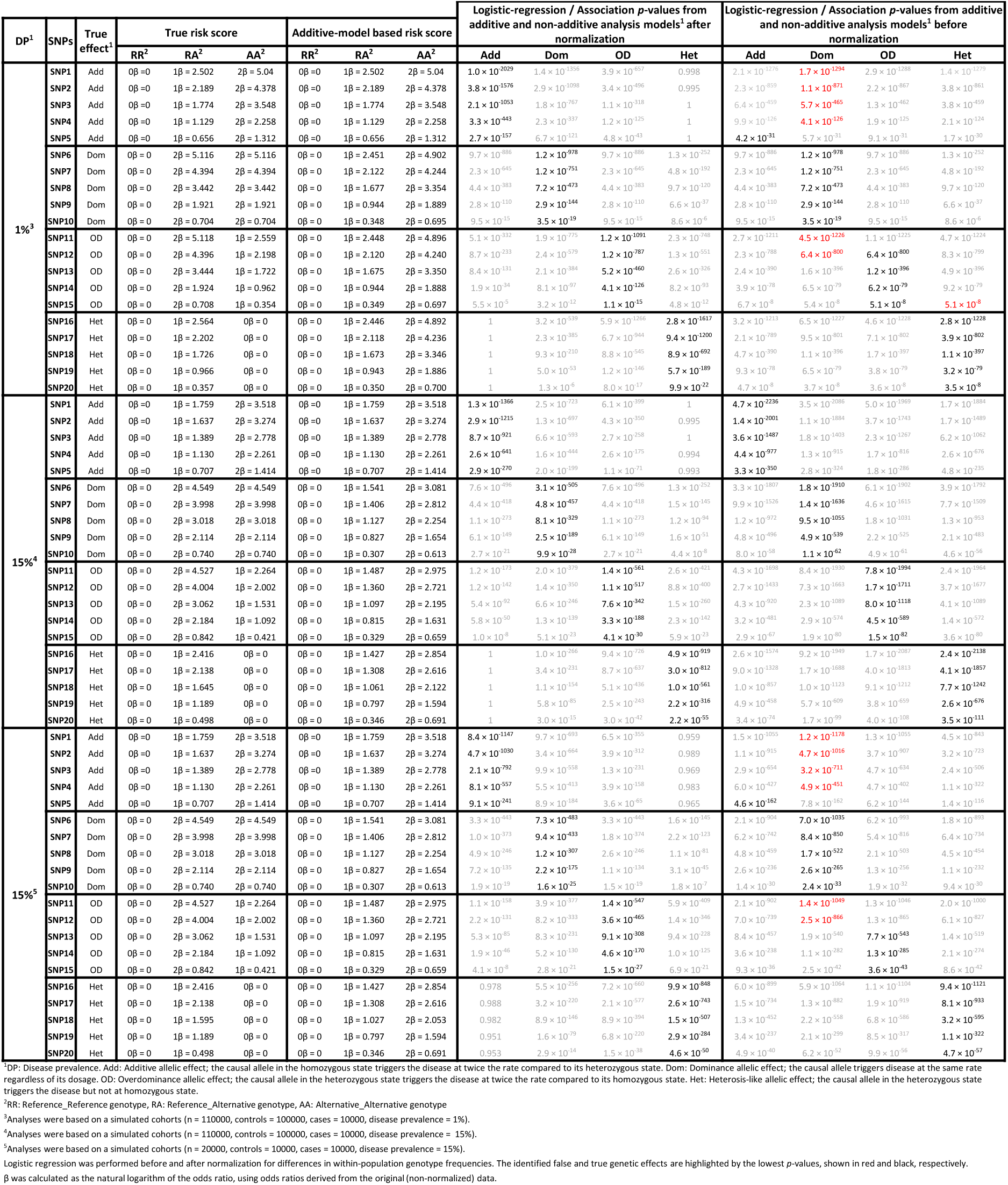
Additive-model based association analyses tend to produce biased risk-score estimates under conditions of non-additive allelic interactions and a pre-normalization step is needed to account for the differences in within-population genotype frequencies and thereby, for the detection of genetic effects through regression line fit.

**Table S2.**
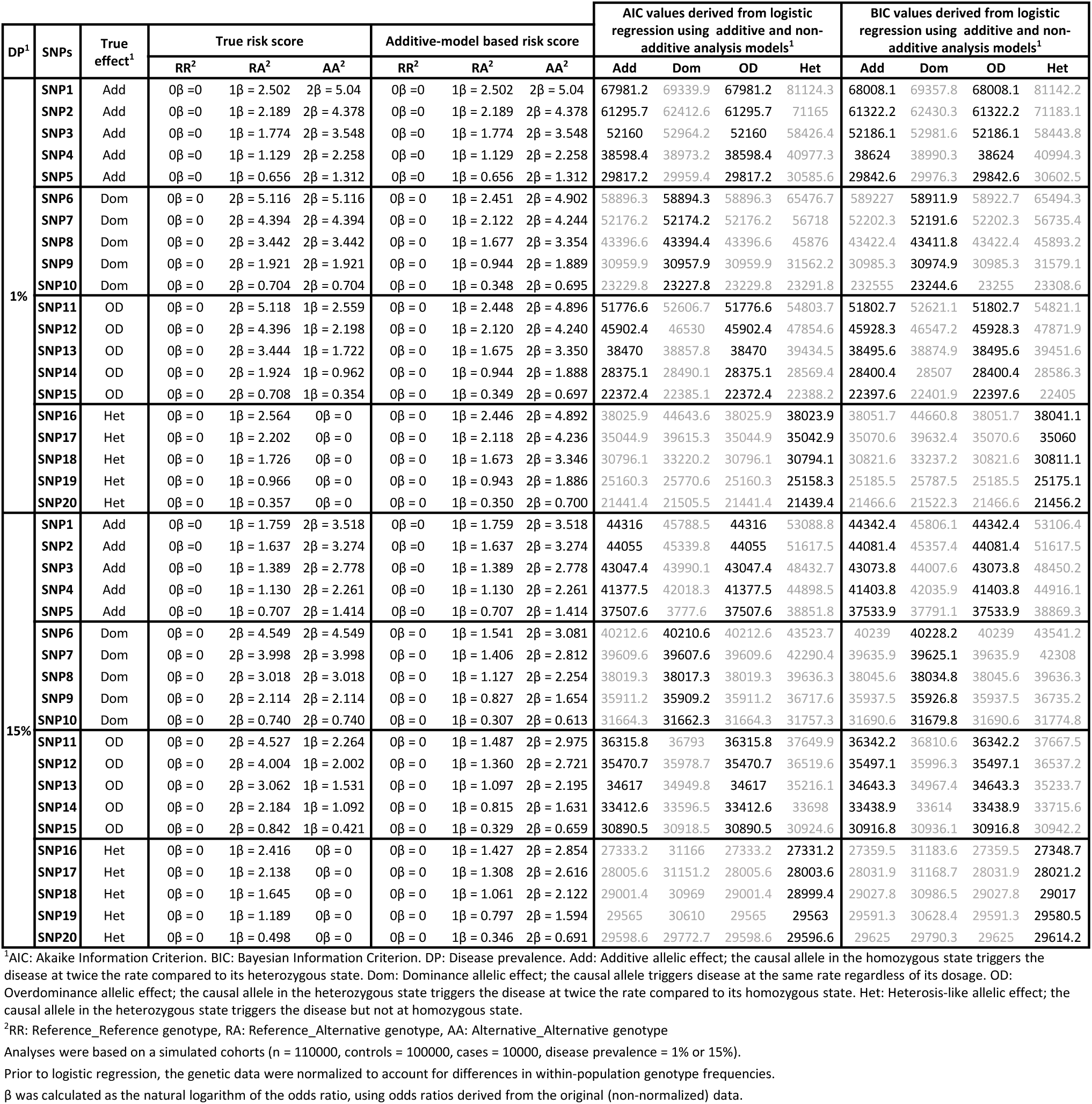
The Akaike Information Criterion (AIC) and Bayesian Information Criterion (BIC) were not capable of distinguishing between additive and overdominance effects.

**Table S3.**
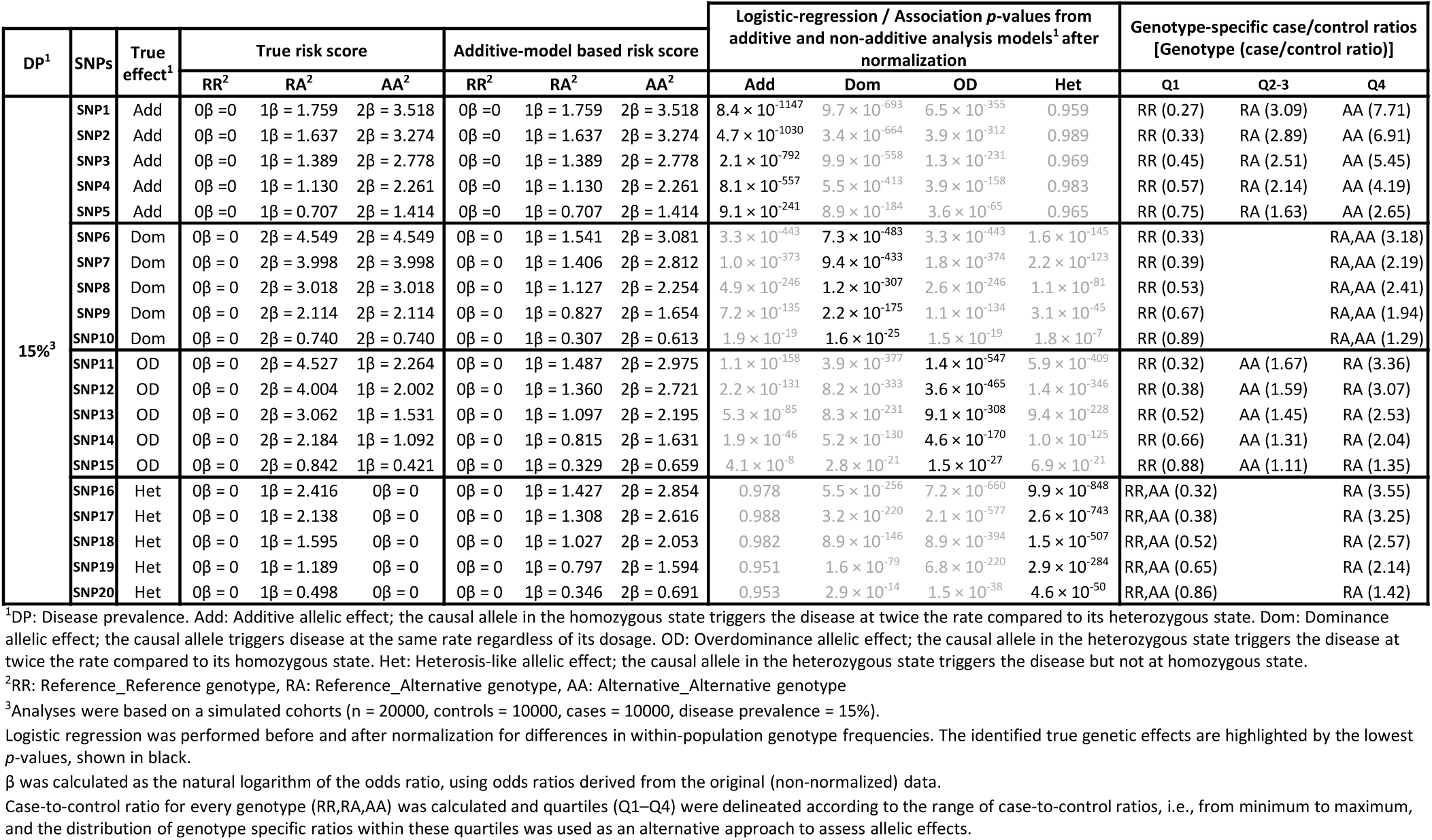
Additive-model based association analyses tend to produce biased risk-score estimates under conditions of non-additive allelic interactions.

